# The importance of reaction norms in dietary restriction and ageing research

**DOI:** 10.1101/2022.12.23.521778

**Authors:** Mirre J P Simons, Adam J Dobson

## Abstract

Ageing research has progressed rapidly through our ability to modulate the ageing process. Pharmacological and dietary treatments can increase lifespan and have been instrumental in our understanding of the mechanisms of ageing. Recently, several studies have reported genetic variance in response to these anti-ageing interventions, questioning their universal application and making a case for personalised medicine in our field. As an extension of these findings the response to dietary restriction was found to not be repeatable when the same genetic mouse lines were retested. We show here that this effect is more widespread with the response to dietary restriction also showing low repeatability across genetic lines in the fly (*Drosophila melanogaster*). We further argue that variation in reaction norms, the relationship between dose and response, can explain such conflicting findings in our field. We simulate genetic variance in reaction norms and show that such variation can: 1) lead to over- or under-estimation of treatment responses, 2) dampen the response measured if a genetically heterogeneous population is studied, and 3) illustrate that genotype-by-dose-by-environment interactions can lead to low repeatability of DR and potentially other anti-ageing interventions. We suggest that putting experimental biology and personalised geroscience in a reaction norm framework will aid progress in ageing research.

## Diet and Ageing

Dietary restriction (DR) potently and arguably universally extends lifespan across species (Fontana & Partridge, 2015; Nakagawa et al., 2012; Simons et al., 2013). Various methods that restrict food intake or uptake lead to an extension to lifespan (Taormina & Mirisola, 2014). These pro-longevity effects were initially interpreted as a beneficial effect of slowing metabolism through reduced caloric intake (Masoro, 2005; Redman et al., 2018). However experiments varying components of the diet, separating mainly macronutrients, suggested that calories alone did not explain the life-extending effects of these treatments (Grandison, Piper, et al., 2009; Mair et al., 2005; Min & Tatar, 2006). By varying carbohydrate and protein content in a geometric framework (Simpson & Raubenheimer, 1993), restriction of dietary protein was suggested as a key determinant of longevity for insects and mice (Piper et al., 2011; Solon-Biet et al., 2014). However, whether dietary protein alone, more specifically amino-acid availability, is directly responsible for DR’s pro-longevity effect has recently been brought into question by experiments in the fly (*Drosophila melanogaster)* (Gautrey & Simons, 2022; Zanco et al., 2021). Furthermore, the effects of calories have been suggested to be more important than considered in the geometric framework in mice (Speakman et al., 2016). Despite a large increase in our knowledge of how diet and especially its restriction affects lifespan we still have limited certainty of which components of the diet, and which associated physiology are responsible for these pro-health effects.

An additional complication to our understanding of ageing is that responses to DR and pro-longevity drugs vary genetically (Jin et al., 2020; Liao et al., 2010; McCracken, Adams, et al., 2020; McCracken, Buckle, et al., 2020; Rohde et al., 2021; Swindell, 2012; Unnikrishnan et al., 2021; Zhou et al., 2014). If this genetic variation is substantial, it offers therapeutic promise for personalising dietary and anti-ageing treatments to increase their effectiveness and translational potential (M. B. Lee & Kaeberlein, 2018; Perez-Matos & Mair, 2020; Sierra et al., 2021). To be able to apply and fully understand how anti-ageing treatments work, we must therefore separate genetic effects from dietary and environmental effects. An important additional complication here is that differences in dietary composition, level of DR or dose of anti-ageing drugs can interact with both environmental and genetic effects (David et al., 2005; Garte, 2006). Some genotypes could have a different dose to longevity relationship that could depend on environmental conditions as well. Such genetic variation in the dose-response to a treatment or the environment is termed a reaction norm in ecology and evolution (Nussey et al., 2007). Reaction norms have important implications for how genetic variance in response to treatments are interpreted (Flatt, 2014; Tatar, 2011).

### Implications of a reaction norm perspective for dietary restriction and ageing research

Here, we will discuss and demonstrate three important implications of a reaction norm perspective on DR and ageing research. 1) Variation in reaction norms can be over- or under-estimated if only a limited number of doses are tested. 2) When a population of heterogeneous genetic makeup is tested, this generates a population-level reaction norm that is a composite across many reaction norms, and this can skew results. 3) Genotype-by-dose-by-environment interactions can lead to low repeatability of DR and potentially other anti-ageing interventions, explaining inconsistencies and equivocations in our field.

### What are reaction norms?

In ecological and evolutionary research reaction norms are used as a term to describe the genetically encoded plastic response to the environment (Flatt, 2014; Nussey et al., 2007). The appreciation of reaction norms stems from an interest in phenotypic plasticity; the plastic nature of organismal responses to the environment (West-Eberhard, 1989), which is present even in organisms devoid of genetic variation, such as clones. Examples for which reaction norms are evident are behaviour and growth (Giebelhausen & Lampert, 2001; Schlichting & Pigliucci, 1995). The incorporation of reaction norms in quantitative genetic statistical models has shown genetic variance in reaction norms to the environment (Brommer et al., 2005; Nussey et al., 2005; Strickland et al., 2021). Furthermore, its importance in evolution is exemplified by the finding of different reaction norms between populations of closely related species (Murren et al., 2014). In medicine, or more precisely pharmacology, dose-response curves (including pharmacodynamics) could be seen as analogous to the foundation of maximising treatment efficacy whilst avoiding harm in the terms of unwanted side-effects (Gabrielsson et al., 2010). Dose-response curves and reaction norms thus describe fundamentally similar processes.

Reaction norms for phenotypes across an environment tend to take a concave shape, with a peak or optimal response observable. Such a concave shape (Figure 1) can represent a wide range of traits and treatments. For diet, DR is said to be occurring where lifespan is maximised and it is at this diet where there is often also a suppression of reproduction observed (Chippindale et al., 1993; Moatt et al., 2016). We further know that if we impose too much restriction, we see starvation or malnutrition and a reduction in lifespan. At fully fed conditions, maximum performance, or optimal Darwinian fitness at which organisms reproduce most offspring during their lifetime is observed (Jensen et al., 2015). If we go beyond this point and overfeed an organism, we see obesity or toxicity effects (McCracken, Buckle, et al., 2020). Non-linear reactions with nutrition are found in humans as well, in which the relationship between nutritional metrics and biomarkers of health often follow a concave relationship (Senior et al., 2022).

**Figure 1.**
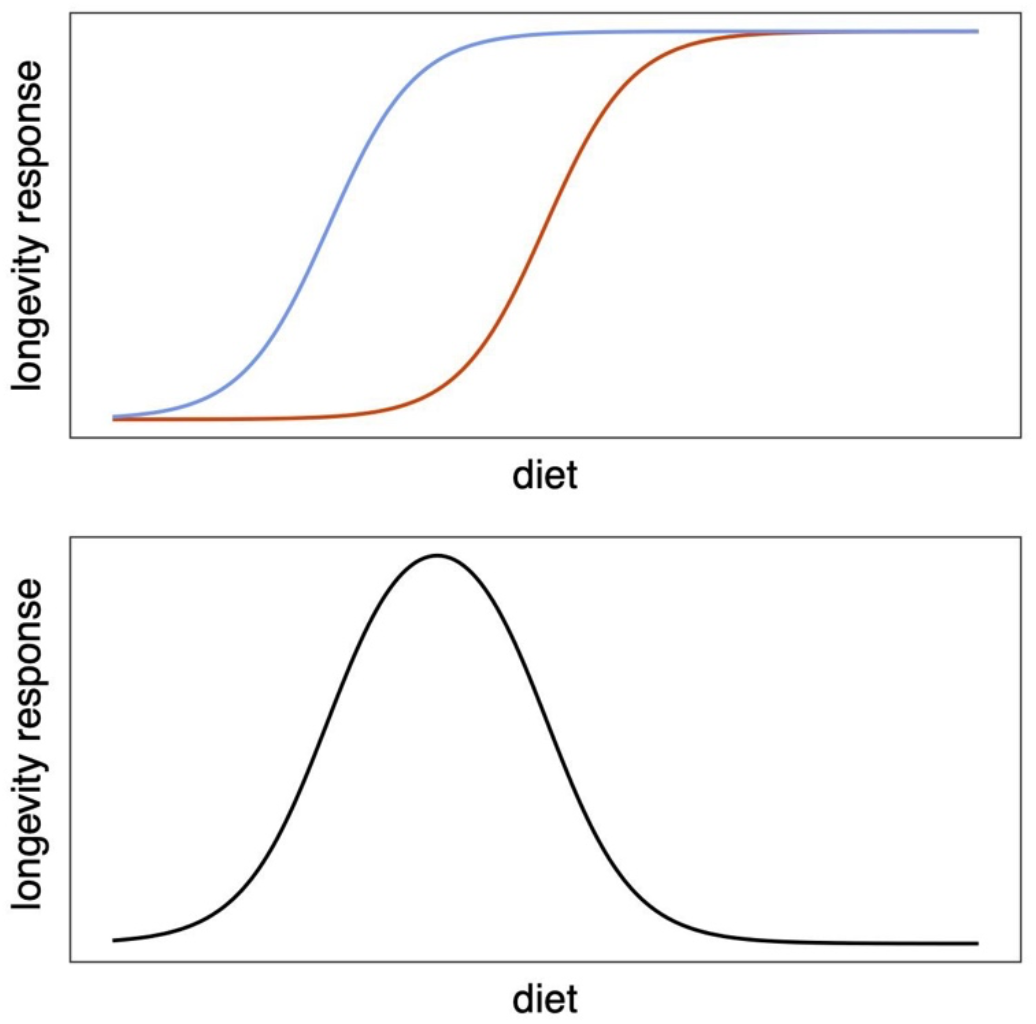
Example of the reaction norm concept. The difference between a benefit (blue) and cost (red) logistic function when shifted in nutritional space yields a concave reaction norm. The same genotype will respond to diet according to this reaction norm. This framework can be expanded to include more nutrients with differential cost functions which in concert determine the overall phenotypic response. Importantly these costs and benefit functions need not be similar in shape and relative importance across all genotypes and environmental contexts.

We propose that such a concave shape arises from the intersection of reaction norms for cost and benefit. In pharmacology, concentration responses are most commonly described by a logistic shape, i.e. S(sigmoid)-shaped curve, with slow increases in response at low doses, before eventual saturation at high doses (Finney, 2009). The ideas of trade-offs between costs and benefits (Stearns, 1989; Winder et al., 2022) are common in biology and in ageing research (Cohen et al., 2020; Kirkwood & Austad, 2000), with the optimum investment in a certain trait or process traded off against its cost. In the specific case of diet, the phenotypic benefits of a given level of consumption will be determined by the underlying physiological costs and benefits. In this framework, we note that the difference between two S-curves, one depicting benefits and one harm, provides a concave curve of the sort that is commonly observed phenotypically (Figure 1). In other words, the phenotypic response to dietary variation will be straightforwardly determined as physiological benefits of a given diet, minus physiological costs. The optimum diet for a particular trait, such as lifespan, is therefore the one on which the difference between the sigmoids for cost and benefit is maximised. It is therefore intuitive to think of reaction norms as a composite of many dose-responses that need not have the same scales or shapes. In biology and medicine, we are often concerned with the optimisation of all such costs and benefits and hence a concave reaction norm framework is intuitive.

Changes in the shape of reaction norms can be categorised in two main ways, a shift in the x-or y-plane but retaining the same shape, and a change in the amplitude (and/or shape) of the reaction (Figure 2). These two changes are often conflated in interpretation, with conclusions made about shape or shift in the response without the accompanying evidence. Unfortunately, it is challenging to conduct experiments on sufficient scale for resolution to discriminate between these two possibilities. Nevertheless, such interpretation is important, as it implicates how physiology is altered. For example, if the DR response is absent due to a reduction in the sensitivity to malnutrition (the left-hand side of the reaction norm) this is a different interpretation than suggesting the anti-ageing mechanisms of DR are affected by a manipulation. In general, when there is a shift in reaction norm but not shape, a change in the set-point response to the environmental condition is more probable, with downstream physiology left intact.

**Figure 2.**
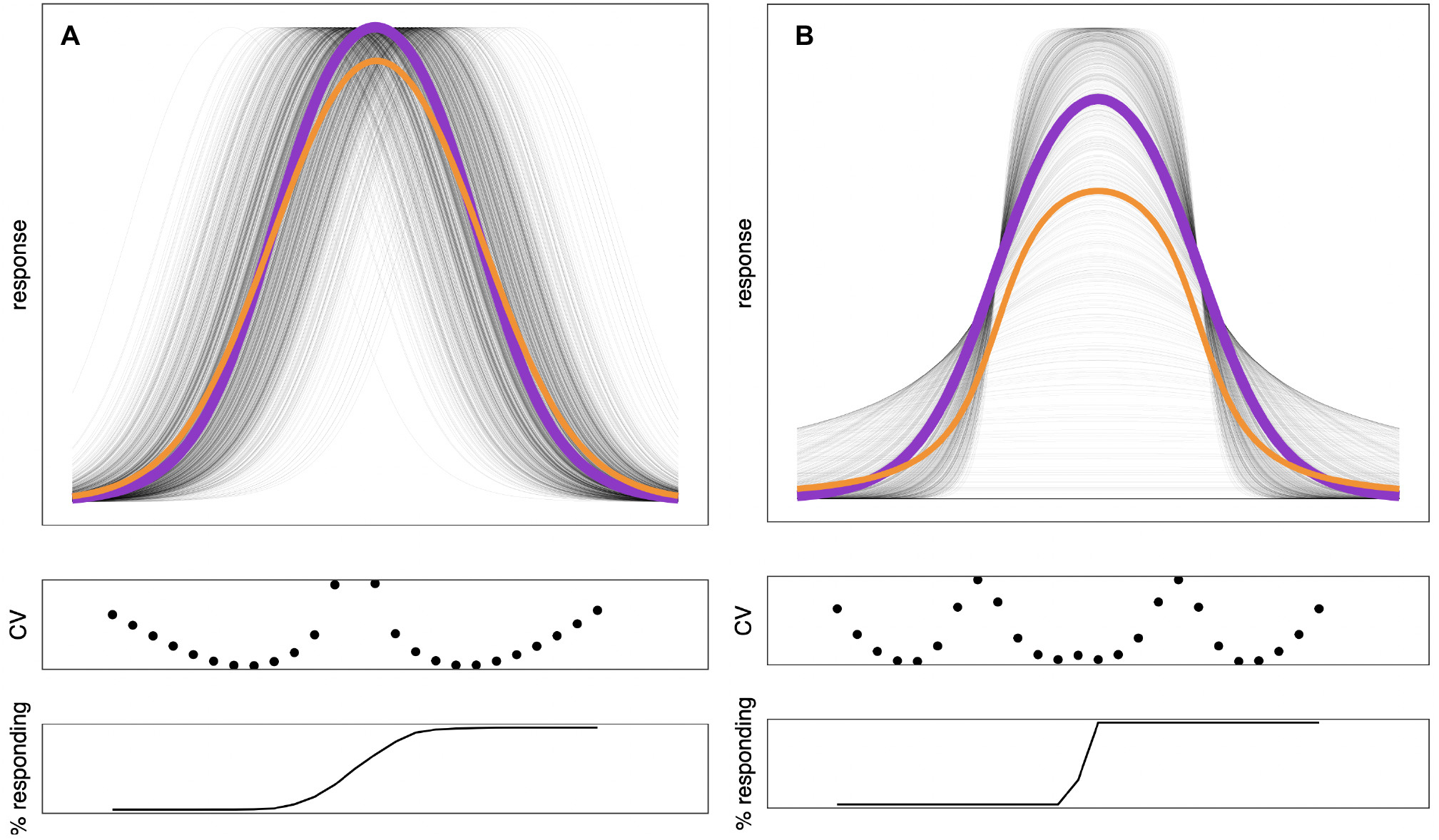
A simulation of genetic variance (normally distributed) in location (A) and shape (B) of the reaction norm. A thousand reaction norms were generated and are plotted across as thin black lines. The purple thick line indicates the genotypic mean simulated with the orange indicates the average response when taken as an average across the phenotypic variance generated. The coefficient of variance (CV) is given as an estimate of variance in the phenotypic space assessed by using a dyad of diets around that location in the reaction norm. Similarly, the number of individuals assigned as a dichotomy between responders or non-responders is given.

Indeed, experiments with mutants have claimed to identify genes involved in the DR response, but a shift in the reaction norm curve is often observed, rather than a change in shape (Tatar, 2007, 2011). These observations indicate that these mutants may not mediate the downstream physiology responsible for the anti-ageing effects of DR, but rather change the point at which DR becomes apparent. Genetics of the response to DR can also be studied in naturally varying populations, although titrations of a range of diets have rarely been applied in multiple populations. The limited studies available suggest that standing genetic variation in the lifespan reaction norm to DR can be found in both a change in both location and shape of the curve (Grandison, Wong, et al., 2009; McCracken, Buckle, et al., 2020; Metaxakis & Partridge, 2013; Zhu et al., 2014). However, comparisons among these studies are challenging, as effects are often not expressed in comparable manner, i.e. in a relevant effect size such as hazard ratio (Gautrey & Simons, 2022; McCracken, Buckle, et al., 2020).

### Variation in reaction norms and their interpretation

We argue that studies of dietary titration in different genotypes will be necessary to comprehensively understand how DR works. Most investigation to date has however been limited to *Drosophila*. Studies in flies have shown genetic variation for reaction norms of lifespan to diet (Grandison, Wong, et al., 2009; McCracken, Buckle, et al., 2020; Metaxakis & Partridge, 2013; Zhu et al., 2014). Studies in other species are so far limited to investigation of how two strains of mice responded to two levels of restriction (Mitchell et al., 2016), and studies in yeast, in which a range of media conditions can be tested (Schleit et al., 2013). It is understandable that studies using a range of diets are limited in number as they require an increasing amount of effort with an increasing number of diets tested. A decision is therefore often made to test dyads of diets across more genetic lines (Jin et al., 2020; Liao et al., 2010; McCracken, Adams, et al., 2020) compared to testing multiple diets across a more limited amount of lines. Conversely, most studies using a wider range of diets often rely on testing one genotype, and extremes of this approach can be found in geometric framework studies (Jensen et al., 2015; K. P. Lee et al., 2008; Skorupa et al., 2008; Solon-Biet et al., 2014).

That genetic variation in lifespan reaction norms to diet is relatively understudied makes it possible that genetic variation in lifespan reaction norms to diet could be even larger than we are currently expecting based on these limited studies. Similar arguments can be made for differences in response to nutrition between the biological sexes as for genotypes. The inclusion of both sexes in nutritional and ageing research is of recognised but still of underappreciated importance (Chen et al., 2022; Garratt, 2020). The dogma in the field to date has been that females are more responsive to DR than males. However, if we recast this in a reaction norm framework, we note that absence of the response in one sex at the same dyad of diets does not necessarily mean the one sex is refractory to the response (e.g. Regan et al., 2016), as their reaction norm might simply be shifted. Arguably we therefore need a dose response curve within each sex to determine how the physiological reaction to nutrition and ageing is changed by biological sex. In a pharmacological context differences in response to anti-ageing treatment could be explained by bioavailability and/or receptor density rather than a difference in the downstream physiology (Garratt, 2020), further increasing complexity of the interpretation between the sexes that would also apply across genotypes.

### How measurement resolution of reaction norms affects dietary restriction and ageing research

When assessing genetic variance to DR it is common to use a dietary dyad (Jin et al., 2020; Liao et al., 2010; Unnikrishnan et al., 2021). Although understandable, this strategy does come at a particular cost, however, as it could lead to falsely concluding DR does not extend lifespan in some genotypes. An absence of a DR effect can be due to a shift in the reaction norm, rather than a change in shape. Moreover, interpretation mistakes can be made suggesting that there is genetic variance for the DR longevity response, whereas there could, for instance, be a change in the susceptibility to malnutrition only. When genetic variance is assessed more generally the position in the reaction norm at which phenotypes are assessed will determine the magnitude of genetic variance in the response measured. The DR response can thus be over- or under-estimated if only a limited number of doses or diets are tested, and failure to detect a DR response may reflect genetic variance in reaction norms. To assess the extent of these biases we simulated how the magnitude of genetic variance is dependent on the location and shape of the lifespan reaction norm to DR.

The estimates of genetic variance from a dyad of diets were highest around the peak of the reaction norm curve when there is a shift in the reaction norm (Figure 2A). In contrast, a change in the shape of the reaction norm resulted in estimates of variance to be highest on the inflection points of the reaction norm curve (Figure 2B). Importantly, when data is analysed as a dichotomy between responding and non-responding lines, a shift in the reaction norm flattens the area in which non-responding genotypes are found. These results are intuitive but not obvious when interpreting variation in responses to DR. Moreover, they imply that genetic variance in reaction norms could affect interpretation in important ways.

Science is iterative and DR is optimised through successive studies or titration within one genotype. Therefore, the level of DR imposed on animals is most likely that of the average genotype in the population. Thus, variance in the reaction norm across other genotypes will be estimated around the peak of the modal reaction norm, as it is there that the strongest DR response is observed for the average genotype. It is also at that point in the reaction norm that most variation in the response can be observed (Figure 2A) and where the dividing line between estimated responders and non-responders is located (Figure 2). When, for example, recombinant inbred lines (Liao et al., 2010) or inbred isofemale lines (Jin et al., 2020) are used to generate genetic variation, there is a risk that phenotypes are assessed around the region of peak DR. A large fraction, or even half of the genotypes will then be pushed into malnutrition and will thus be designated as non- or low responders. Indeed, prior studies on mice and flies, which compared longevity between two diets, found a surprising fraction of genetic lines that lacked a response to DR (Jin et al., 2020; Liao et al., 2010).

### Could genotype-by-reaction norm interactions reduce repeatability?

Our model predicts that subtle differences between laboratories could change set-points and shapes of response to DR such that a dichotomous conclusion of “did / did not respond” may not accurately represent the complexity at play, and effects that may have been observed on a different region of a diet dilution may be missed. Thus, a genotype that responds to a dyad DR intervention in one lab may fail to do so in another, because of small differences in diet (Piper et al., 2014) or genotype affecting reaction norms, and repeating the dyad therefore misses the “right” region of the response curve. Indeed, the genetic variance in the longevity effects induced by DR have low repeatability although up until now data were only available from a partial repeat of DR experiments in recombinant inbred lines of mice. When the same mouse strains (Liao et al., 2010) were tested again later, findings were not replicated (Unnikrishnan et al., 2021).

We report here now that in flies, a similar pattern can be observed across a wide range of genotypes. We compared published data using a dyad of diets in the Drosophila Genetic Reference Panel (DGRP, Mackay et al., 2012) to unpublished data from the Simons laboratory (Figure 3). Several aspects of the experimental setup between these two studies differ. Jin *et al*. housed flies in vials, whereas Simons used cages, and there are small differences in diet (Simons diets as in McCracken, Adams, et al., 2020), especially in relative sugar and cornmeal content. Sample size was similar (N = 100 for Simons, and N = 200 for Jin *et* al.). In the literature there is one more study (Dick et al., 2011) testing DR across the DGRP but they housed flies as mixed sex, with fewer lines tested, and therefore we did not include this study in our comparison. Within each study lifespan between diets correlated across the genetic lines tested (*r*_s_ = 0.56, p < 0.001; Jin, r_s_ = 0.33, p < 0.01; Simons). Across studies lifespan was correlated weakly under fully fed (r_s_ = 0.20, p = 0.08) and DR (r_s_ = 0.24, p = 0.04) conditions. When both conditions were averaged per study and consequently correlated a moderate and significant correlation could be observed (r_s_ = 0.32, p < 0.01). Part of the genetic effect determining lifespan across the genetic lines tested was therefore replicable. In contrast, the magnitude of the DR response (difference in lifespan between DR and fully fed condition) was not replicable across labs (r_s_ = 0.15, p = 0.21). Despite this, the large majority of genetic lines showed a longer lifespan under DR in both labs (52 out of the 72 lines included, p < 0.01 compared to 50% responding).

**Figure 3.**
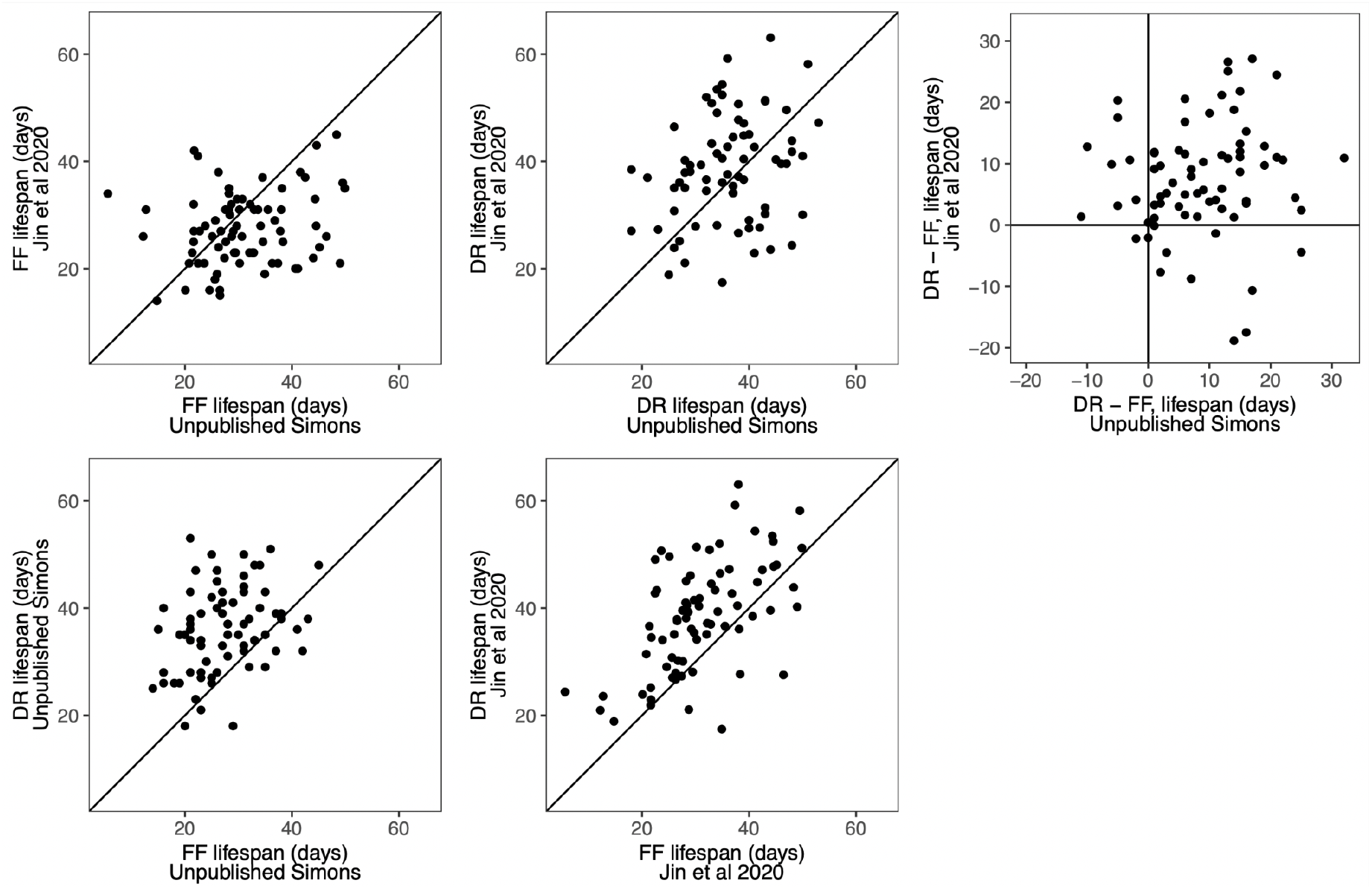
The lifespan of the same genetic lines of flies measured in two laboratories under DR and fully fed conditions. Lifespan was moderately repeatable within (bottom panels) and weakly repeatable across (top left two panels) studies at both diets, but the relative response to DR showed low non-significant repeatability (top right corner).

It appears therefore that the DR longevity response is not commonly repeatable. We suggest that such low repeatability is most probably explained by small differences in the food, the lab environment or their interaction inducing small shifts in the reaction norm. Such small changes in reaction norms can dramatically change the response to two diets. Fitting with this observation, gene by environment interactions for lifespan are readily observed even in the lab environment (Leips & Mackay, 2000). When diet is manipulated many such environmental interactions are manipulated in multivariate nutritional space. Left largely underappreciated in DR research especially is that single nutrients within this space can interact and could determine longevity in specific circumstances only. Thus, observations of specific nutritional components explaining DR (Gautrey & Simons, 2022; Grandison, Piper, et al., 2009; Zanco et al., 2021) could be relevant in one context only. As such the reaction norm between diet and longevity (Figure 1) should be seen as a concert of different interacting nutrients that shift in importance according to environmental and genetic effects. An important implication of this is that certain genetic backgrounds might be predisposed to certain ageing trajectories depending on the environmental conditions at which lifespan or phenotypic ageing is assessed.

### Population level genetic variance and reaction norms

Small genetic differences in the genetic lines between the copies of strains kept by different labs induced through genetic drift, selection or bottlenecks could explain why lifespan itself was not highly repeatable (Figure 3). As such it is hard to generate a strategy to effectively assess variation to the DR longevity response apart from assessing complete reaction norms to diets across a limited number of lines. It has been suggested that using outbred strains is a solution to this problem as it reflects a mixture that arguably is also more representative of a natural population (Grandison, Wong, et al., 2009; Mair et al., 2005; Sgrò et al., 2000). However, the mean reaction measured need not be the same as the mean genetic reaction norm of the population. Averaging across phenotypes resulting from genetic variance in both shifts and changes in shape of the reaction norm flattens the overall response measured (Figure 2). The reason for such flattening is that the simulated Gaussian genetic variance in reaction norms results in skewed distributions across points along the reaction norm. Therefore, reaction to diet or any dose-response for that matter is expected to be skewed in outbred populations and not necessarily representative of within-individual reactions. Such effects could be further augmented by frequency dependent selection (Wolf & McNamara, 2012) that is probable to operate in cultures (Kojima & Huang, 1972). As such it could also be unlikely that reaction norms will be stable in such populations as they will be subject to drift and selection (Selman & Swindell, 2018). Subsampling from these populations could also result in reduced repeatability.

An elegant solution to inherently genetically unstable outbred populations is to cross two or more inbred stocks together to generate a heterogenous population of offspring for experiments of known and largely stable genetic ancestry. The HET3 mice used in highly controlled trials for anti-ageing interventions are a good example of this strategy (Harrison et al., 2019; Miller et al., 2002; Selman & Swindell, 2018). In such a strategy it is arguably less important to understand or estimate the genetic variance in reaction norm. If the intervention and dosage does not work well for the majority of the population, a response would not be detected and is thus not relevant to move into human clinical trials. However, a strong assumption here is that the genetic variance present in the model study population is genetic variance relevant for humans. Any such reasoning rests on the assumption that the specific lines used harbour genetic variants that are present across species and that these variants have similar effects across species. Currently we have limited evidence that genetic associations found across species are doing so through the same conserved loci, most probably because a lot of genetic variance is based on polymorphisms in regulatory regions which are divergent between species (Flint & Mackay, 2009). Perhaps most importantly we know that species have widely divergent standing genetic variation (Gossmann et al., 2012). On a qualitative level we thus do not know whether there is the same genetic variance in model organisms for the physiology of the trait or reaction norm of interest. On a quantitative level we do not know whether the level of genetic variance is proportionally similar to humans. It therefore remains unclear whether an absence or presence of an effect in a model organism is a good predictor for universal replication in other model organisms and most importantly humans.

## Conclusions

A reaction norm perspective on ageing research and nutrition is important to not mistake variation in reaction norms for a *de facto* absence of an effect. This argument has been made before (Flatt, 2014; Tatar, 2007, 2011; Voelkl & Würbel, 2021), but is worth reiterating especially with conflicting data emerging in our field without reference to the reaction norm framework. In addition, we expand this framework here to include genetic variation within the population of study and show relatively low repeatability of DR-induced longevity within flies. Hidden variation in reaction norms within a study population can reduce the overall amplitude of the reaction detected in phenotypic space. We argue that this hidden variation could lead to the low repeatability of DR’s longevity benefits that have been reported and we report here. Note that these reasons will also apply to any other dose-response relationship, and we expect are not exclusive to diet. It is perhaps surprising that quantitative models (Hadfield & Kruuk, 2010) incorporating genetic relatedness (Brommer et al., 2005; Nussey et al., 2005) have not been used to study genetic variance in dose responses in laboratory populations (Bou Sleiman et al., 2022). Having a population level estimate of genetic variance in the reaction norm will quantify the confounding effect reaction norms impose on experimental biology in ageing research. Conversely, personalised applications of geroscience (M. B. Lee & Kaeberlein, 2018; Perez-Matos & Mair, 2020; Sierra et al., 2021) will require the identification of individuals that benefit from certain anti-ageing treatments. Understanding such personalisation in a dose-response, i.e. reaction norm framework, should aid progress in the specific and growing area of personalised geroscience.

## Acknowledgements

MJPS is supported by a Sir Henry Dale Fellowship (Wellcome and Royal Society; 216405/Z/19/Z) and an Academy of Medical Sciences Springboard Award (the Wellcome Trust, the Government Department of Business, Energy and Industrial Strategy (BEIS), the British Heart Foundation and Diabetes UK; SBF004\1085). AJD is supported by a UKRI Future Leaders Fellowship (MR/S033939/1), the Biotechnology and Biological Sciences Research Council (BB/W510658/1), and a University of Glasgow Lord Kelvin Adam Smith Fellowship. For the purpose of Open Access, the author has applied a CC BY public copyright licence to any Author Accepted Manuscript version arising from this submission. We thank Richard Kaskiewicz, Laura Hartshorne, Gracie Adams and Andy McCracken for helping to collect the unpublished data included in this manuscript.

## Declarations

The authors declare no competing interests.

